# Single-neuron representations of spatial targets in humans

**DOI:** 10.1101/523753

**Authors:** Melina Tsitsiklis, Jonathan Miller, Salman E. Qasim, Cory S. Inman, Robert E. Gross, Jon T. Willie, Elliot H. Smith, Sameer A. Sheth, Catherine A. Schevon, Michael R. Sperling, Ashwini Sharan, Joel M. Stein, Joshua Jacobs

**Affiliations:** Doctoral Program in Neurobiology and Behavior, Columbia University, NY, NY 10027, USA; Department of Biomedical Engineering, Columbia University, NY, NY 10027, USA; Emory University School of Medicine, Atlanta, GA 30322, USA; Department of Neurosurgery, University of Utah, Salt Lake City, UT 84112, USA; Department of Neurosurgery, Baylor College of Medicine, Houston, TX 77030, USA; Department of Neurology, Columbia University Medical Center, NY, NY 10032, USA; Department of Neurology, Thomas Jefferson University, Philadelphia, PA 19107, USA; Department of Neurosurgery, Thomas Jefferson University, Philadelphia, PA 19107, USA; Department of Radiology, Hospital of the University of Pennsylvania, Philadelphia, PA 19104, USA

## Abstract

The hippocampus and surrounding medial-temporal-lobe (MTL) structures are critical for both memory and spatial navigation, but we do not fully understand the neuronal representations used to support these behaviors. Much research has examined how the MTL neurally represents spatial information, such as with “place cells” that represent the current location or “head-direction cells” that code for the current heading. In addition to behaviors that require an animal to attend to the current spatial location, navigating to remote destinations is a common part of daily life. To examine the neural basis of these behaviors we recorded single-neuron activity from neurosurgical patients playing Treasure Hunt, a virtual-reality spatial-memory task. By analyzing how the activity of these neurons related to behavior in Treasure Hunt, we found that the firing rates of many MTL neurons during navigation significantly changed depending on the position of the current spatial target. In addition, we observed neurons whose firing rates during navigation were tuned to specific heading directions in the environment, and others whose activity changed depending on the timing within the trial. By showing that neurons in our task represent remote locations rather than the subject’s own position, our results suggest that the human MTL can represent remote spatial information according to task demands.

## Introduction

The medial temporal lobe (MTL) is critical for memory and spatial navigation (Scoville and Milner, 1957; Morris et al., 1982). Many electrophysiological studies have focused on characterizing MTL neuronal coding during navigation, and much of this interest in spatial navigation is due to the fact that the neuronal mechanisms underlying spatial coding are thought to relate to those used for memory (Eichenbaum, 2017; Miller et al., 2013; Schiller et al., 2015; Epstein et al., 2017). Place cells, whose firing rates change as a function of an animal’s location in space, are arguably the most well studied cell type in the MTL (O’Keefe and Dostrovsky, 1971), and show activity related to navigation and mnemonic processing (Miller et al., 2013). Similarly, the MTL contains other neurons that activate according to an animal’s spatial setting, such as grid and head-direction cells (Hafting et al., 2005; Jacobs et al., 2013; Taube et al., 1990), which could also have broader functional roles Bellmund et al. (2016); Horner et al. (2016). Building off this literature, a topic of growing interest is whether the types of coding patterns that represent an animal’s own spatial location are also used to represent other kinds of information to support complex behaviors, such as the targeting of remote locations during goal-directed navigation (Pfeiffer and Foster, 2013; Vass et al., 2016).

In addition to thinking about one’s own location, everyday life often involves remote locations— in particular, planning, remembering, and navigating to remote destinations. However, the neuronal representations of remote locations remain less well understood compared to those of an animal’s current location (Spiers and Maguire, 2007). Beginning to address this, a growing line of studies examined how MTL neurons represent salient remote locations (Rolls and O’Mara, 1995; Wilming et al., 2018; Wirth et al., 2017; Omer et al., 2018; Danjo et al., 2018). There is also evidence that certain single-neurons and hippocampal-BOLD signals activate to represent particular views and goals during navigation to specific fixed navigational goals (Ekstrom et al., 2003; Ainge et al., 2007; Brown et al., 2016). This diverse literature indicates that the MTL can represent various aspects of an animal’s current spatial context according to behavioral demands.

We wanted to go beyond earlier studies that probed memory for locations marked by visual landmarks (eg. Miller et al. (2013); Wirth et al. (2017)) and investigate more generally how the human MTL codes for current remote spatial target locations. To examine this issue, we asked neurosurgical patients with microelectrodes implanted in their MTL to play a virtual-reality spatial-memory task and we examined how their neural responses related to their simultaneous movement. The fifteen participants in our study played Treasure Hunt, a video-game–like memory task that measured subjects’ ability to remember the spatial locations where various objects were hidden (Miller et al., 2018). In each trial of the task, subjects explored a virtual beach and traveled to a series of treasure chests. Upon reaching each chest, it opened and an object appeared. The subject’s goal was to encode the spatial location corresponding to the position of each item. In contrast to earlier tasks with human single-neuron recordings that tested memory for fixed spatial landmarks, objects in Treasure Hunt were placed at previously unmarked locations in an open environment, allowing us to identify whether the brain utilizes a neural coding pattern for remote locations that is related to the one used by place cells during movement. Our primary result is identifying “spatial-target cells,” which are neurons whose firing rates were modulated according to the location of the remote target, rather than the subject’s own location. We also identified neurons that altered their firing rate according to the subject’s heading, and others whose firing rate was modulated as a function of the timing within each trial. The range of responses we observed—including the representation of remote goals by spatial-target cells—indicate that the human MTL represents multiple types of spatiotemporal context information to support goal-directed spatial processing.

## Results

To examine the neural signatures of navigating to and remembering remote destinations, we asked fifteen neurosurgical patients with implanted microelectrodes implanted to play Treasure Hunt, a virtual-reality spatial-memory task. In Treasure Hunt subjects navigate across a rectangular environment using a handheld joystick and learn the locations of objects hidden at each of various spatial locations Miller et al. (2018). In each trial of the task subjects learn two or three trial-unique objects that are each positioned at random locations (see Methods). During the task, we recorded the activity of 131 neurons (Fried et al., 1999), from the hippocampal formation (HF; n=45), entorhinal cortex (EC; n=71), and parahippocampal cortex or perirhinal cortex (PC; n=15). To identify the neural patterns that support behavior this task, we tested whether neuronal firing rates during navigation were significantly modulated by the locations of upcoming to-be-remembered objects as well as the subject’s current location, heading direction, timing within the trial, and subsequent memory performance.

Each trial of Treasure Hunt consisted of three phases (Fig. 1a; see Methods). First, the subject navigated to a chest (“Navigation”). Upon reaching a chest, it opened and was either empty or revealed an object whose location the subject was instructed to remember (“Encoding”). In a given trial subjects repeated this for up to four chests. Next, in the “Recall” phase subjects were asked to recall the object locations. Each subject in our study performed one of the two task versions, which differed solely in terms of the Recall phase. In the “object-cued” task version, subjects viewed the image of a cued object and used the joystick to move an on-screen cursor to indicate its remembered location. In the “location-cued” version of the task they viewed a probed location and verbally responded by speaking the name of the corresponding object into a microphone that recorded their response.

**Figure 1:**
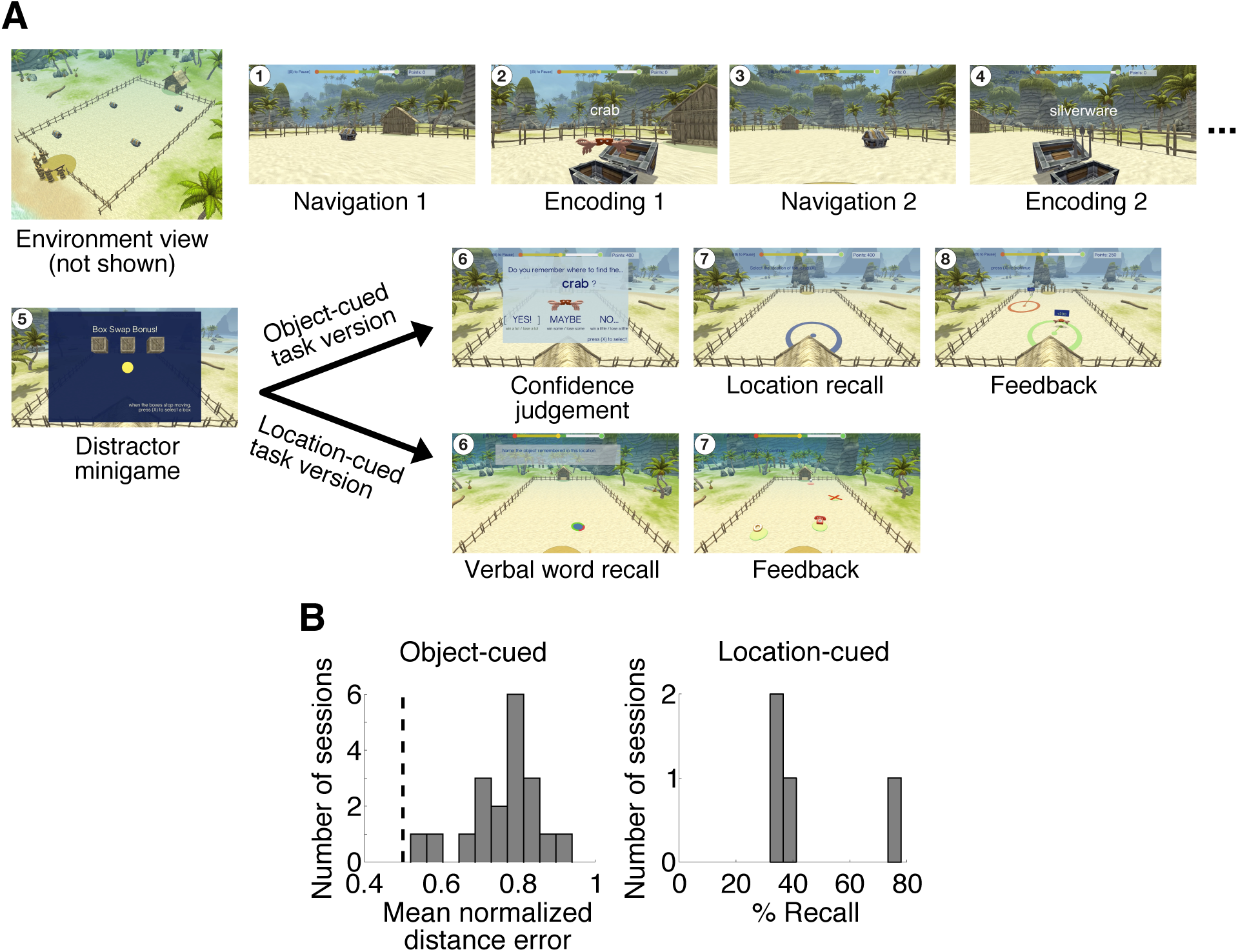
Behavior in the Treasure Hunt task. A. Timeline of Navigation, Encoding, and Recall events in Treasure Hunt. Numbered panels indicate the sequence of events that subjects encounter in each trial of the task. The object- and location-cued task versions differ in terms of the recall stage, which is indicated here via the divergence at Step 6 in this timeline. See Methods for details on the structure of the two task versions. B. Histogram showing mean performance on the recall phase of Treasure Hunt in each of the two task versions. Left, Distribution of mean normalized distance errors across sessions for the object-cued task version. A normalized distance error of 1 corresponds to the best possible response, and 0.5 corresponds to chance performance. Right, distribution of mean percentage of items that were vocalized correctly in the location-cued task version.

### Subject behavior in Treasure Hunt

We assessed task performance by scoring the subject’s performance during Recall for each object–location pair. For the object-cued task version we assessed accuracy for each studied item by computing the distance between the subject’s response location and the item’s actual position, and then converting this distance error to a accuracy measure that accounted for all possible distance errors (see Methods). The left panel of Figure 1b shows a histogram of the mean accuracy, or normalized distance error, in each object-cued task session (N=19 sessions). Mean accuracy across all object-cued sessions was 76.7 ± 2.2%, and the performance in all individual sessions were above chance (0.5). The recall phase of the location-cued Treasure Hunt task is similar conceptually to commonly used paired-associate memory tasks. The right panel of Figure 1b shows a histogram of each location-cued session’s mean percent recall (N=4 sessions). The mean performance on location-cued sessions (46%) was consistent with levels seen in other paired-associate memory tasks that required verbal responses (e.g., Greenberg et al. (2015)).

We found that subjects maintain, and, in fact slightly improve, memory performance throughout each session. On average, in the object-cued task version subjects’ memory performance is 6.6% better in the second half of each session compared to the first half (p=0.026; paired t-test), which demonstrates that subjects successfully learn to perform the task better within a single session. We also examined how memory performance varied according to the order of an object–location pair within each trial. Figure S1 shows the mean accuracy for presented objects as a function of the position within the trial for all object-cued sessions. There was a significant effect of serial position on memory performance (*p* = 0.008; repeated measures ANOVA), likely reflecting improved memory for items at primacy and recency positions, consistent with patterns seen in conventional verbal memory tasks (Kahana, 2012).

### Neurons responsive to the current spatial target

We were interested in whether neurons represented information about the location of the current, to-be-remembered chest during navigation. We were motivated by previous work in animal models showing MTL cells that represent salient remote locations (Rolls and O’Mara, 1995; Wilming et al., 2018; Wirth et al., 2017; Omer et al., 2018; Danjo et al., 2018; Gauthier and Tank, 2018), as well as related evidence from recordings of human theta oscillations (Lee et al., 2018) and fMRI (Brown et al., 2016). Therefore, we examined how the firing rates of individual neurons during navigation varied according to the location of the current target chest.

To identify these “spatial-target cells,” we analyzed each cell’s spiking activity as a function of the location of the upcoming chest, in addition to the subject’s current position (Fig. 2a). We generated spatial maps of each cell’s firing patterns, based both on the location of the upcoming spatial target and on the subject’s own position. We then identified neurons whose firing rates were significantly modulated by these factors using a permutation procedure based on an ANOVA. We labeled a neuron as a spatial-target cell if its firing rate significantly varied as a function of the upcoming chest location at *p* < 0.05. Because chests had the same appearance and objects were not visible during navigation, spatial target cell firing must reflect information about the spatial location of the upcoming target. As an example of this phenomenon, Figure 2b illustrates the activity of one example spatial-target cell from the left entorhinal cortex of Patient 9. This neuron increased its firing rate when the subject navigated to chests that were located in the “south-central” part of the environment (*p* < 0.001).

**Figure 2:**
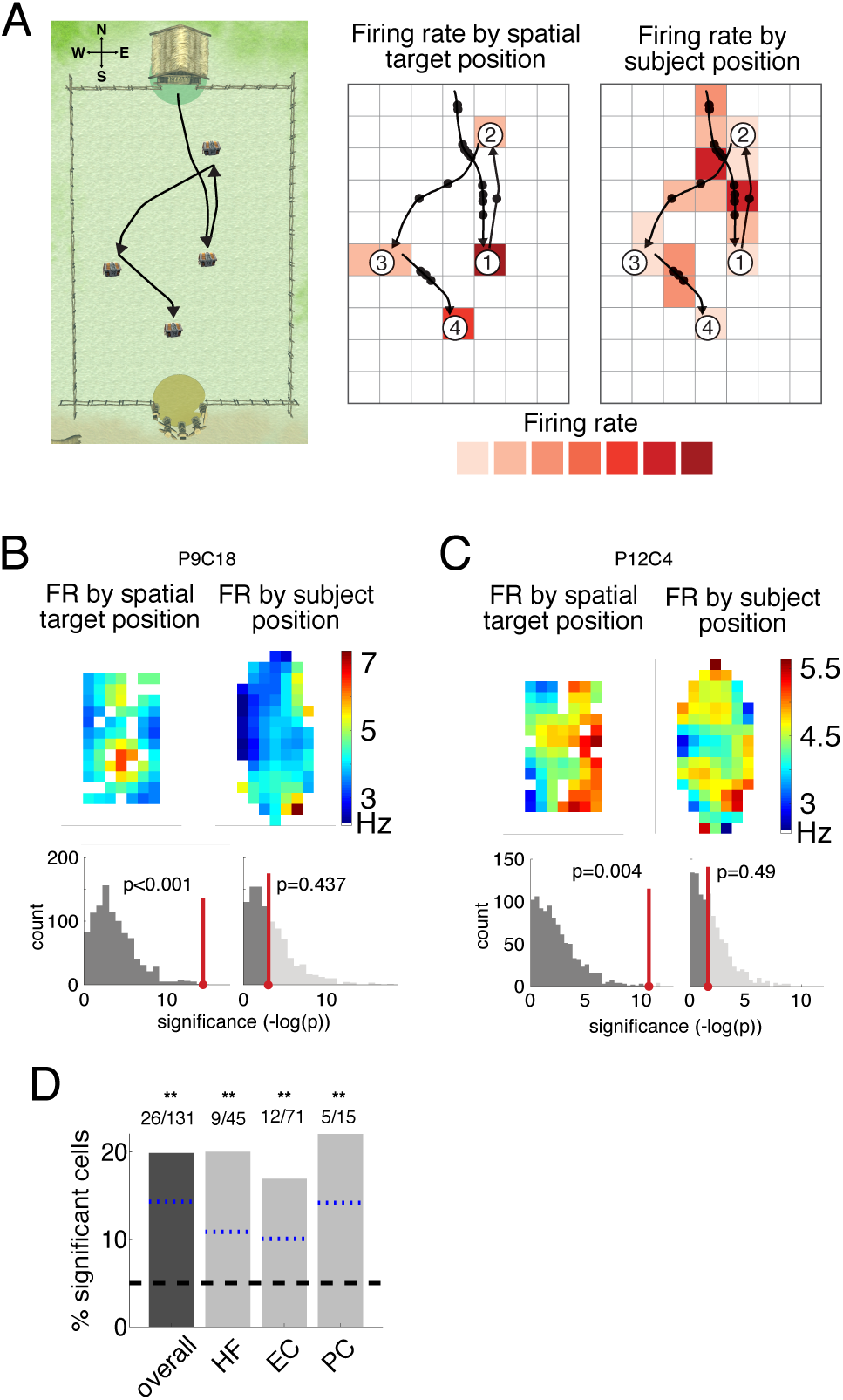
Neural activity related to spatial target position. A. Analysis framework for binning navigation period neuronal data by subject position and spatial target location, shown for an example trial. Left, overhead view of Treasure Hunt environment with example paths to 4 chests (only one chest is visible at a given time). The NESW coordinates we use are shown in the upper left. Middle, example path spikes binned by spatial target location to calculate firing rate during navigation based on the chest location. Right, same spikes binned by subject position to calculate firing rate on the path. B. Top-left, firing rate map of navigation activity binned by spatial target position for a neuron in the left entorhinal cortex from Patient 9. Black line indicates the perimeter of the traversable virtual environment, and areas that didn’t meet minimum traverse time requirements as described in the Methods are plotted in white.. Bottom-left, histogram of p-values from ANOVA (see Methods) assessing spatial target location modulation of firing rate for the observed data (red) versus shuffled data (gray). This cell’s activity is significantly modulated by the spatial target position (permutation-corrected ANOVA, p < 0.001). Top-right, firing rate map for current location. Bottom-right, histogram of p-values from ANOVA assessing current location modulation of firing rate. Neuron is not significantly modulated by subject position (p = 0.43). C. Same as B but for another example neuron in the right entorhinal cortex from Patient 12. Neuron is significantly modulated by spatial target position (p = 0.004) and not subject position (p = 0.49). D. Percentage of significant spatial-target cells by region. Shown for all MTL neurons (“overall”) and also split into HF, EC, and PC. Symbols above the bars indicate p-values from a one-sided binomial test for each proportion, FDR-corrected (* * p < 0.01, *p < 0.05, +p < 0.1). The black dashed line represents the 5% false positive rate, and the blue dotted lines are the lower 95% confidence interval from the one-sided binomial test for each bar.

Critically, while this example cell’s firing rate was modulated by the location of the upcoming chest, its firing rate did not vary significantly according to the subject’s own position (Fig. 2b, right panel; *p* = 0.4). Figure 2c shows a second example of this phenomenon from a cell in Patient 12’s right entorhinal cortex. This cell significantly increased its firing rate when the subject approached spatial targets in the “east” section of the environment (*p* = 0.004), and also did not show a firing-rate modulation according to the subject’s own position (*p* = 0.4). Because the firing of these cells was modulated by the location of the upcoming to-be-remembered target and not the subject’s own position, the activity of these neurons constitutes a novel coding pattern that is distinct from the activity of conventional place cells.

To assess the statistical reliability of this phenomenon, we first confirmed that the number of identified spatial-target cells was significantly greater than expected by chance, followed by correcting for multiple comparisons across the five behavioral variables we examined (see Methods). Across the population, 20% of MTL cells (26 of 131) were classified as significant spatial-target cells. This proportion was significantly more than the 5% expected by chance (*p* = 8.3×10^−9^, one-sided binomial test, FDR-corrected). The number of spatial-target cells was also significantly above chance when measured separately for the hippocampal formation, entorhinal cortex, and PC regions (one-sided binomial tests *p*’s= 5 × 10^−4^, 5 × 10^−4^, 6.1 × 10^−4^ respectively, FDR-corrected). We found significant spatial-target cells in eleven of the fifteen subjects who participated in our study. The prevalence of spatial target cells did not significantly differ between task versions (see Supplemental Text).

We next examined the timecourse of spatial-target cell activity. Figure S7b–d plots the activity of the spatial-target cell from Figure 2a over time, split by paths to a location in versus out of the cell’s firing field. The results of this analysis at the group level (Figure S7e) show that spatial-target cells exhibit elevated firing throughout navigation periods to chests in their firing field, indicating that the activity of these cells is not related to transient behavioral events at the beginning or end of each navigation epoch. To examine the possibility that spatial-target cells reflected information about the distance to the upcoming target, we tested for distance-related activity during navigation. Six spatial target cells also showed distance-related activity, but this overlap between the two cell types was not significant (p=0.12, *χ*^2^ test, df=1). Furthermore, to rule out the possibility that spatial target cell responses could be explained by navigation time, for each spatial-target cell we compared the durations of the navigation periods leading to chests inside versus outside of the cell’s firing fields using a two-sample t-test. At the group level we did not find that path duration for navigation to chests in and out of the spatial-target cells’ firing fields differed for a significant proportion of cells (2/26, p=0.4, one-sided binomial test). Overall, these results further support our finding that a substantial number of neurons throughout the human MTL specifically represent remote locations in our task.

### Analysis of neuronal activity related to the current spatial location

In addition to representing the location of the current chest, an additional potentially relevant spatial variable is the subject’s own location during navigation. In light of the extensive literature on spatial coding for self location in various species (O’Keefe and Dostrovsky, 1971; Ekstrom et al., 2003), we tested for neurons in this dataset whose firing rates were modulated by the subject’s own position. Using the ANOVA described in the preceding section, we identified the neurons whose firing rates were significantly modulated as a function of the subject’s own position. Figure S2 shows three example “place-like” cells that showed spatial modulation according to the subject’s current virtual location at a level of *p* < 0.05 when measured individually (see also Figure S8a.) However, at the population level the proportion of place-like cells was not significantly greater than the 5% expected by chance after multiple comparison correction (8 of 131 = 6%; *p* = 0.33, one-sided binomial test, FDR-corrected; Fig. S2d). As such, during the navigation phase of Treasure Hunt we do not find strong evidence that the firing rates of individual MTL neurons are correlated with the subject’s own location.

### Neurons responsive to heading direction

In rodents there is evidence for neurons whose firing rates are modulated by the direction of the animal’s head during movement (Taube et al., 1990; Robertson et al., 1999). These head-direction cells are commonly found in the dorsal presubiculum and anterodorsal thalamus, but have also been found in areas of the MTL such as the entorhinal cortex (Sargolini et al., 2006). These cells have not previously been found in humans and this gap is notable because a neuronal representation of orientation is important for navigating to a new location in an environment. We therefore tested for the existence of “heading-modulated” cells in our dataset, which we defined as neurons that varied their firing rate according to the direction that subjects moved in the virtual environment. Figure 3a–d illustrates the activity of four significant heading-modulated cells. As these examples illustrate, individual heading-modulated cells showed peak firing activity at differing headings. In addition, some cells showed increased firing rates at multiple distinct headings (Figure 3c–d), similar to “bidirectional cells” observed in rodents (Jacob et al., 2017).

**Figure 3:**
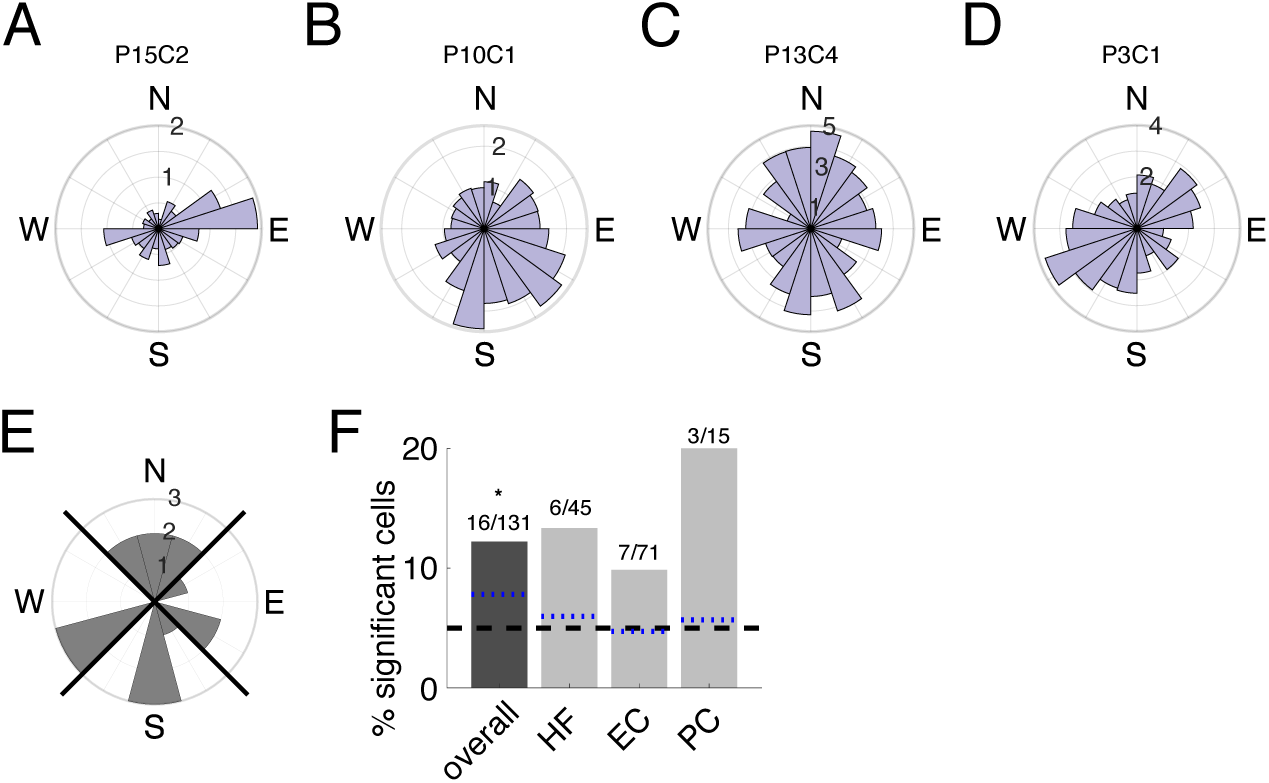
Neural activity related to the subject’s heading. Firing rate by virtual heading direction (N=“north”, E=“east”, S=“south”, W=“west”) for example cells significantly modulated by heading direction. A) Circular histogram of firing rate by heading direction for example cell from Patient 15, in the left hippocampus, significantly modulated by heading direction (p = 0.026). Firing rate is indicated with numbers on the concentric circles. B. Example cell from Patient 10, in the left parahippocampal cortex, significantly modulated by heading direction (p = 0.005). C-D. Two more significant heading-modulated cells, both in the left hippocampus (from Patient 13; p = 0.0315, from Patient 3; p = 0.003). E. Circular histogram of preferred heading directions for each significant heading direction cell. Counts indicate number of cells with that preferred direction. Black diagonal lines indicate NESW quadrants. F. Percentage of significant heading-modulated cells by region. Symbols above the bars indicate p-values from a one-sided binomial test for each proportion (* * p < 0.01, *p < 0.05, +p < 0.1). The black dashed line represents the 5% false positive rate, and the blue dotted lines are the lower 95% confidence interval from the one-sided binomial test for each bar.

Overall, 12% of MTL cells (16 of 131) showed significant heading direction modulation, which is significantly more than expected by chance (*p* < 0.0015, one-sided binomial test, FDR-corrected). No heading-modulated cells showed firing that significantly varied with the subject’s current position. Three heading-modulated cells showed effects of spatial target position, but this overlap was not significant (*p* = 0.9, *χ*^2^ test). Because the population of heading-modulated cells did not significantly overlap with those that were significantly modulated by spatial target position, it suggests that spatial-target cells are not explained by direction-related modulations, which could have been possible in theory if similar locations were always approached from a particular direction.

Additionally, we did not find that any particular preferred angle was dominant across the population of heading-modulated cells (p=0.97, Rayleigh test; Figure 3e). This was important because due to the trial start locations and the rectangular shape of the environment, “north” and “south” directions were sampled more frequently (Figure S6d), which provided a potential source of bias. Because we did not find evidence of a preferred direction for heading-modulated cells, this suggests that our heading-modulated results are not due to uneven directional sampling.

### Neurons modulated by serial position

In addition to space, a growing body of work shows that neurons in the MTL also represent information about event timing during spatial tasks (MacDonald et al., 2011; Heys and Dombeck, 2018; Tsao et al., 2018). We tested whether MTL neurons in this task represented event timing information by measuring whether neuronal activity during navigation was modulated according to the serial position of each navigation period within each trial.

Subjects navigated to up to four chests in a trial, enabling us to investigate differences in neuronal activity based on the chest order. We analyzed navigation periods based on their serial position, and found that the firing rates of 20 of 131 (15%) of MTL cells were significantly modulated by serial position, which is significantly more than expected by chance (*p* = 2.18×10^−5^, one-sided binomial test, FDR corrected). Figure 4a–b shows examples of two such “serial position” cells. We specifically found a significant proportion (23%) of serial position cells in the entorhinal cortex (Fig. 4d; *p* = 1.05×10^−6^, one-sided binomial test, FDR corrected). Across all cells showing modulation by serial position, there was a preference for these neurons to represent the initial list position (see Figure 4c). The set of serial position cells did not significantly overlap with the spatial target cells (p=0.55; *χ*^2^ test, df=1). Five serial position cells were also classified as heading-modulated cells, which is notable but not a statistically significant overlap (p=0.058; *χ*^2^ test, df=1).

**Figure 4:**
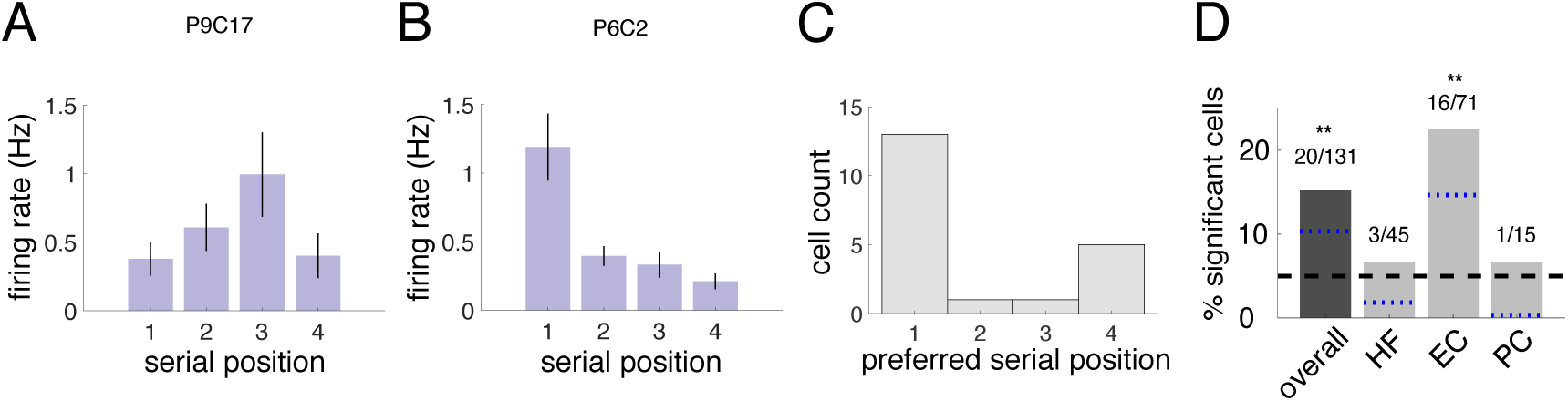
Neural activity related to serial position. Firing rate by serial position for two example cells significantly modulated by the serial position of the chest. Error bars are SEM of the trial means in each condition. A. Example neuron from Patient 9 in the left EC, significantly modulated by serial position (p = 0.025). B. Example neuron from Patient 6 in the right EC, p < 0.001. C. Histogram of preferred serial position for each significant serial-position cell. Counts indicate number of cells with maximal firing rate in that serial position. D. Percentage of significant serial position cells by region. Symbols above the bars indicate p-values from a one-sided binomial test for each proportion (* * p < 0.01, *p < 0.05, +p < 0.1). The black dashed line represents the 5% false positive rate, and the blue dotted lines are the lower 95% confidence interval from the one-sided binomial test for each bar.

### Analysis of neuronal activity modulated by subsequent memory

Studies in humans with large-scale neural activity such as functional MRI and local field potentials often find that neural activity in the hippocampal formation during encoding increases in relation to subsequent memory performance (Schacter and Wagner, 1999; Miller et al., 2018). However, little evidence for this finding exists at the single-neuron level for changes in mean firing rates (Rutishauser et al., 2010). We therefore tested whether each cell’s firing rate during navigation significantly varied as a function of whether or not the current spatial target was subsequently remembered. We found eleven “memory cells”, whose firing rate during navigation varied as a function of whether or not the current spatial target was subsequently remembered (see Fig. S3a–b). This constitutes 8% of cells, which is not significantly greater than the 5% expected by chance (*p* = 0.081, one-sided binomial test, FDR-corrected). Thus, our results do not provide strong evidence for a population of MTL neurons whose mean firing rates during navigation correlate with subsequent memory performance (see Supplemental Text for more information).

## Discussion

In this study we found that during a spatial-memory task the firing rates of subjects’ MTL neurons were significantly modulated by the locations of spatial targets, heading direction, and serial position. In particular, by showing that human single-neurons can represent information about remote spatial positions, these results help explain how contextual information, such as relevant remote locations, are represented by the brain to support goal-directed spatial navigation and cognition.

The spatial-target cells that we identified share some properties with MTL neurons reported in previous human and animal studies. Namely, Ekstrom et al. (2003) identified cells in the human MTL that activated during navigation to specific fixed navigational goals (“goal cells”). Broadly, the activity of both goal cells and spatial-target cells could be interpreted as related because they both code for aspects of a subject’s current objective during navigation. However, as we describe below, we believe that the distinctive features of our task allowed us to differentiate between spatial-target cells and goal cells in at least three key ways. First, owing to the pseudo-random placement of chests in our task, we were able to determine that spatial-target cell firing is directly linked to specific locations in the virtual environment. In contrast, Ekstrom et al.’s goal cells may respond according to the landmark at a particular location, as opposed to the location itself (see also Komorowski et al. 2009). Second, spatial-target cells likely support a more location-focused type of goal coding, which we were able to identify due to the greater path variability enabled by our open-field environment. In contrast, it is likely that Ekstrom et al.’s goal cells reflect route–goal conjunctions because their task required that subjects follow stereotyped routes along constrained paths to fixed landmarks, which is correlated with route–goal conjunctive representations in rodents (Kobayashi et al., 2003; Grieves et al., 2016). Third, there is substantial evidence that spatial tuning and hippocampal activity differ dramatically according to task type (Markus et al., 1995) and memory-state (Wood et al., 2000), and, specifically, between memory and navigation states (Miller et al., 2018). Since our task differed from Ekstrom et al.’s in all three aspects by requiring memory encoding, it is likely that spatial-target cells reflected a distinct kind of goal coding compared to Ekstrom’s goal cells, which showed this pattern during navigation.

Ekstrom et al. (2003) also identified “view cells,” which, along with the “spatial view cells” identified by Rolls in nonhuman primates, are neurons that fire in response to a subject viewing specific objects or locations. However, both of these cell types differ in important ways compared to the spatial-target cells we report. Whereas Ekstrom et al.’s view cells respond to a visual stimulus’s presence anywhere on the screen, the spatial-target cells we describe instead activate according to the precise location of the to-be-remembered target in the spatial environment. Further, Rolls’ spatial-view cells differ from our results by reflecting a different kind of spatial representational scheme. Spatial-view cells showed increased activity when the animal viewed locations in the context of a particular 2-D scene (usually spots on the wall of a room). In contrast, because our task allowed subjects to approach individual target locations from drastically different headings, it allowed us to conclude that the activity of spatial-target cells reflected navigable locations within open 3-D space, irrespective of the background scene. Nonetheless, despite these substantial apparent differences between our results and the earlier findings, it remains possible that there are important links between all these cell types. Understanding the representational similarities across these cell types is an interesting area of future work that could be accomplished with specifically targeted behavioral tasks.

Other studies have also shown MTL place-like cells whose firing patterns are modulated by information related to the current navigational goal (Frank et al., 2000; Ainge et al., 2007; Ferbinteanu and Shapiro, 2003; Wirth et al., 2017), which is conceptually related to our finding of human spatial-target cells. However, the results from those studies differ significantly from ours because they show goal-related neural activity that appeared only at particular locations along a track, which were often near choice points or goal locations. Broadly, this is an important difference because spatial-target cells seem to be involved in continuous encoding of the target location, while the goal-related place cells described above are likely involved in context-based decision making en-route to goals.

In addition to identifying neurons that activate for remote target locations, our finding of heading-modulated cells provides perhaps the first evidence of this cell type in humans (see also Jacobs et al. 2010). Head-direction cells have been described extensively in rodents and are most frequently found in areas such as the postsubiculum, retrosplenial cortex, anterodorsal thalamus (Taube et al., 1990; Taube, 1998; Robertson et al., 1999). However, they have also been found in the hippocampus and entorhinal cortex (Leutgeb et al., 2000; Sargolini et al., 2006). In addition to the cells we found that respond most to movement in a single direction, we also found evidence of cells with bidirectional responses, which are similar to patterns reported recently in rodents (Jacob et al., 2017). Additionally, although none of the heading-modulated cells we report were localized to the subiculum, the bidirectional heading-modulated cells also bear similarities to the subiculum cells described in Olson et al. (2017) that were tuned to opposing directions. An interesting area of future work will be to determine the degree to which these cells are similar, especially given the differences in the navigational setting between our task and those in the other studies.

Finally, it is notable that we also found neurons that activated to represent particular serial positions during each trial, and that these cells were prominent in the entorhinal cortex. Because this population of neurons showing “serial position” effects did not significantly overlap with the neurons showing “spatial target” or memory effects, it suggests that these phenomena reflect distinct neural processes. An interesting area for future work is identifying the extent to which serial-position cells are relevant for understanding other aspects of memory. The localization of serial-position cells to the entorhinal cortex aligns with recent work in rodents and human fMRI that identified a specific role for the entorhinal cortex in supporting the neural encoding of time through an experience (Tsao et al., 2018; Montchal et al., 2019). Further, serial-position cells most often showed increased activity during navigation periods at the beginning of each trial (Fig. 4c), which is notable because entorhinal time cells also show distinctive responses at early moments in a sequence. This correspondence suggests that these responses may be related (Tsao et al., 2018; Bright et al., 2019).

The nature of MTL spatial coding can vary with task demands (e.g., Aronov et al. 2017). We believe the lack of observed place cells in our dataset may be related to the behavioral demands of our task. In our experiment, subjects tried to remember the positions of objects that had been placed at locations in an open environment that were subsequently unmarked. On each trial of our task, new, previously unmarked locations become salient and relevant for memory encoding. This may have led to greater attentional focus to those upcoming locations during navigation instead of the subject’s current location. This focus on remote locations during navigation is a substantial difference between our task and many previous neural studies of human spatial navigation (e.g., Jacobs et al. 2013) and we hypothesize that this feature of our task may have been important for eliciting the activity of spatial-target cells. An interesting direction for future research will be to determine the degree to which the activities of task-related neurons are modulated according to current task demands. This would require a task in which the relevant behavioral factors, such as the type of memory content to be encoded or retrieved, is changed across trials while the same neurons are recorded. This kind of experiment could show whether individual MTL neurons alter the nature of their information coding depending on the features of the environment or current behavioral demands.

Our results extend the prior literature by demonstrating that MTL neurons are engaged during the encoding of remote spatial locations in allocentric space. We also find coding of heading direction, and timing within each trial. A key direction for future work in this area is understanding the degree to which the activities of these cell types are maintained across different behavioral settings. Systematically characterizing the activity patterns of these neurons across behaviors can be challenging, especially in clinical recording environments. To this end it will likely be useful to utilize more advanced measures of behavior, such as incorporating eye tracking and using tasks with multiple conditions that distinguish neural signals related to memory and other behaviors. An additional important direction going forward is to identify links between the multiple different types of neurons that represent task-relevant information in a given setting, such as by identifying relations between place and spatial-target cells. Identifying these links could open directions for future research on what causes these representational schemes to change and show how the brain links the representations of related memories.

## Methods

### Experimental Task

The subjects in our study were neurosurgical patients who voluntarily performed our Treasure Hunt spatial-memory task in free time between clinical procedures. Treasure Hunt is a 3D virtual spatial-memory paradigm developed in Unity3D, which we previously used to study various aspects of human spatial memory and electrophysiology (Miller et al., 2018; Maidenbaum et al., 2019b,a).

Subjects played Treasure Hunt on a bedside laptop computer and controlled their movement through the virtual environment with a handheld joystick. Each patient performed one of two versions of Treasure Hunt (referred to as ‘object-cued’ and ‘location-cued’ in Figure 1a). (For more details on this task version, see Miller et al. (2018).) We included data from both task versions to increase statistical power, but our conclusions remain robust when only the object-cued data are examined (see Supplement). In each trial of Treasure Hunt subjects explored a virtual beach (100 × 70 virtual units) to reach treasure chests that revealed hidden objects, with the goal of encoding the location of each encountered item. The locations of the objects changed across trials, but the environment’s shape, size, and appearance remained constant across the sessions. One virtual unit corresponds to approximately 1 foot in the real world. Subjects viewed the environment from the perspective of biking through the environment and the elevation of their perspective was 5.6 virtual units. As shown in Figure S10, each end of the environment has unique visual cues to help subjects orient. One end of the environment has a beach hut with trees, and the other contains totem poles and a view of the ocean. See the Supplemental Movie for more details on the appearance of the environment.

Each trial of the object-cued task begins with the subject being placed on the ground at a randomly selected end of the environment. The subject then navigates to a chest (i.e., the Navigation phase) using a joystick. (Due to the randomized start locations across trials, the direction of joystick movements are uncorrelated with particular directions in the virtual environment.) Upon arrival at the chest, the chest opens and either reveals an object, which the subject should try to remember, or is empty. The subject remains facing the open chest for 1.5 s (Encoding phase) and then the object and chest disappear, which indicates that the subject should navigate to the next chest that has now appeared in the arena. In each trial the subject navigates to a sequence of four chests. Two or three (randomly selected) of the chests contain an object and the rest are empty. In each session there are a total of 100 full chests and 60 empty chests, over 40 trials. In each trial chests are placed pseudo-randomly across the interior of the environment, subject only to the constraints that no chest can be placed within 11 virtual units from another, and that all chests must be at least 13 virtual units from the arena’s boundary. This 11-virtual-unit restriction ensures that chest locations are varied in a trial and is the cause of the small dip in the center of the occupancy map in Figure S6a–c. There are no constraints based on previous trials, and all object identities are trial-unique and never repeated in a session.

After reaching all four chests of a trial, subjects are transported automatically so that they view the environment from a raised perspective (31 virtual units above ground) at a randomly selected end of the environment. They then perform a distractor task, the “shell game,” before entering the Recall phase. During Recall, subjects are cued with each of the objects from the trial in a random sequence and asked to recall the object’s location. In each recall period, they first indicate their confidence for remembering the object’s location (“high”, “medium”, or “low”). Next, they indicate the object’s precise location by placing a cross-hair at the location in the environment that corresponds to the location of the cue item. After indicating the location of each object from the trial, the Feedback stage of each trial begins. Here, subjects are shown their response for each cued object in the trial, via a green circle if the location was correct and a red circle if it was incorrect. Subjects receive feedback on their performance, following a point system where they receive greater rewards for accurate responses. A response is considered correct if it is within 13 virtual units of the true object location. (Notably, this is a different threshold than the one we use when analyzing subject behavior, described below.)

The location-cued task version is similar to the object-cued task, except that subjects respond differently during the Recall phase. During the Recall phase of the location-cued task, subjects are placed in the same elevated view as in the object-cued version, and view a location cue (a white circle on the floor of the environment). They are asked to respond by verbally recalling the name of the object that was positioned at that location. Each session of this task version consists of 30 trials, each with 3 or 4 chests, for a total of 105 chests per session. None of the chests are empty. During the Recall phase subjects are probed with 4 or 5 locations, one of which is a lure location that does not match the location of any of the trial’s objects. After the Recall phase is complete, subjects receive feedback in the form of the correct objects being placed on top of the circles, and the circle is green if they responded correctly and red if were incorrect. Given our primary interest in characterizing neuronal activity during navigation, for data analyses we pooled from both task versions as they differed only in the Recall phase. An interested reader should see Miller et al. (2018) for information on neural activity during the encoding phase.

### Single-neuron recordings

Fifteen patients (10 male, mean age=32 years, minimum age=20 years) with medication-resistant epilepsy participated in a total of 23 sessions of our task. These subjects were all undergoing a surgical procedure in which depth electrodes were implanted to localize epileptogenic regions. We conducted these recordings at three sites (Columbia University Medical Center, Emory University School of Medicine, Thomas Jefferson University). Electrode placements were determined solely by clinical team. All patients were implanted with Behnke–Fried microelectrodes with 9 platinum–iridium microwires (40 *µ*m) extending from the macroelectrode tip, following previously reported methods (Fried et al., 1999; Misra et al., 2014). All patients provided informed consent. The microwire data were recorded at 30 kHz using NeuroPort recording systems (Blackrock Microsystems, Salt Lake City, UT). Across sessions we successfully isolated 131 putative neurons from microelectrodes in the medial temporal lobe (MTL). Forty-five neurons were in the hippocampus and subiculum (hippocampal formation; HF), 71 were in the entorhinal cortex (EC), and 15 were in parahippocampal or perirhinal cortex (PC) (see Table S1).

### Single-unit waveform classification

We identified neuronal action potentials using the Combinato cluster cutting package (Niediek et al., 2016). Following automatic cluster detection, we manually examined and sorted all clusters. We only included neurons in our analyses after manually inspecting all spike waveforms. This allowed us to visually confirm that all spike waveforms had amplitudes above the noise threshold, physiological-looking waveform shapes, and stationary spiking activity throughout each session. Additionally, we only included units for which more than 95% of spikes had a interspike interval of at least 3 ms, following the criteria from Valdez et al. (2013). We classified any cell as a single unit if its distribution of spike waveform shape fulfilled the criteria from Hill et al. (2011).

We were mainly interested in single-unit activity in this study, and as such we were conservative in our cluster cutting, resulting in 126 of the units being classified as single units and only 5 of the accepted units were classified as multiunits (due to not fulfilling the waveform shape criteria). Across the 131 units the mean percentage of ISIs *<*3ms was 0.27%, with a SEM of 0.03%. Figure S4a shows the waveform of an example unit from Patient 15. For this unit, 0.31% of the ISIs were within 3ms (Figure S4b), and 1.4% of spikes were above the amplitude cutoff (Figure S4c).

### Analysis of behavior in the task

We assessed performance on the object-cued task as in earlier work (Miller et al., 2018). For each response, we defined the distance error as the Euclidean distance between the subject’s response and the correct location. We report ‘accuracy’ as 1 minus the percentile rank of the actual distance error computed relative to the distribution of all possible distance errors that could have been made for the object’s location. This results in an accuracy measure ranging from 0 to 1 (worst to best possible response). We used this normalized metric instead of raw Euclidean distance to separate subjects’ performance from random guessing, since it accounts for the shape and range of possible errors for each response location. Owing to this normalization, random guessing would result in an value of 0.5 on average. We considered an item-location to be correctly recalled if its accuracy was above the session’s median accuracy, and incorrectly recalled if it was below that threshold. Performance on the location-cued task was scored by an experimenter who listened to each recorded audio file manually reported if the spoken word correctly matched the identity of the corresponding memory cue. In the sessions analyzed, subjects never responded with synonyms.

### Electrode localization

Microwire bundle localization followed previously validated protocols (Jacobs et al., 2016; Lee et al., 2018). We determined the anatomical location of each microwire electrode bundle by co-registering the pre-surgical T1-weighted (whole brain coverage, 3D acquisition, 1mm isotropic resolution) and T2-weighted (temporal lobe coverage, coronal turbo-spin-echo acquisition, 0.4 × 0.4 × 2 mm resolution) structural MRIs to the post-surgical CT scan. MTL subregions were automatically labeled using a multi-atlas segmentation technique on the T2-weighted MRI (Wang et al., 2013; Yushkevich et al., 2015). Electrode contact coordinates were then mapped to MRI space and a neuroradiologist (JMS) manually determined the anatomical locations of the microwire electrodes based on the co-registered images (Avants et al., 2008). See Figure S5 for example of the images used to localize microwires.

### Analysis of neural data

Our neural data analyses focused on signals from the navigation periods of each session. We first binned the rectangular environment into a 5 × 7 grid and then computed the grid bin that corresponded to the subject and target locations for each navigation epoch. This 5×7grid provided a level of aggregation for the data from each session, as each grid bin contained data from 10±4 independent chest viewing events (Fig. S6). The spatial target and subject position fell into the same grid location in only 9.5 ± 0.39% (mean ± SEM) of timepoints across the dataset. Next, we measured spatial patterns of neural activity by comparing signals across bins. For the data from a grid bin to be included in the analysis, the subject must have occupied that location for a minimum of 5 s or 2 s when binning by subject position or spatial target position, respectively. We discretized the behavioral navigation data into 100-ms epochs, and for each one calculated the the average x- and y-coordinates and bins for the subject’s location and for the current target chest. We excluded navigation epochs when subjects were still for more than 500 ms. We binned the spike data into matching 100-ms epochs and then calculated the mean firing rates. We smoothed the firing rate maps for visualization purposes by binning into a 11 × 16 grid, applying a Gaussian filter with a 1.1-bin SD, and excluding any grid bins less than 100 ms of occupancy.

We used an ANOVA to identify cells whose firing rates were spatially modulated, as in earlier work (Ekstrom et al., 2003; Miller et al., 2015). Here, the dependent variable was the firing rate of an individual cell in each epoch and the independent variables were the labels of the grid bins corresponding to the subject and target locations. We chose this analysis because it could identify spatial modulation while being agnostic to the size or shape of firing fields and because it could accommodates cell with both high and low baseline firing rates. We felt this flexibility was useful because human place cells sometimes have larger and more complex firing fields compared to neurons recorded in rodents. Further, there is evidence that neurons with complex spatial tuning patterns nonetheless can be used to decode navigationally relevant information that might otherwise be overlooked using traditional procedures (e.g., Hardcastle et al. (2017); Stefanini et al. (2019).

We assessed statistical significance using a permutation test based on circular shuffling, as in earlier work (Ekstrom et al., 2003; Miller et al., 2015). This procedure was repeated 1000 times with circularly time-shifted firing rate values, whereby the firing rate vector for each cell was rotated by a random offset relative to the behavioral navigation data. If the test statistic calculated on the real data was in the 95^th^ percentile of the test statistics from the randomly shifted data, the parameter was considered a significant factor in modulating firing rate. Importantly, the reason we performed this circular shifting of the spike data was to preserve the temporal autocorrelation of the spikes when generating the surrogate data. As a result, the temporal structure of the data was conserved even after shuffling and the resulting p-values from the surrogate distributions were comparable to the observed p-values. It should be understood that this is a conservative procedure, which, in fact, makes it less likely for an observed statistic to be significant. Figure S9 shows the p-values resulting from circularly shifted spikes (panel A) compared to randomly shuffled spikes (panel B) for the example spatial-target cell in Figure 2b.

To test for modulation of firing rate by heading direction, a separate ANOVA was conducted with the heading quadrant (virtual “N”,”E”,”S”,”W”) as factors. The heading directions were determined by measuring the subject’s heading in each epoch of the task and grouping the angles into four 90° bins. We were unable to test the effect of egocentric bearing to the chests on firing rate, because the subject was pointed directly at the chest for the majority of the navigation period. Specifically, during navigation 92 ± 1% of the time subjects were pointed within 60 deg of the chest. To test for modulation of firing rate by subsequent memory we conducted separate ANOVAs with memory performance as a factor. Finally, we tested whether cells’ firing rates were modulated by the serial position. There were 3 or 4 chests in each trial, and we conducted an ANOVA of firing rate by the corresponding serial position of each navigation period.

For each cell-type category, we were interested in whether the observed cell counts significantly exceeded the 5% expected false-positive rate. As such, we tested the observed proportion of significant cells against the null hypothesis that the proportion was less than or equal to chance using one-sided binomial tests with alpha = 0.05. These *p* values were then corrected for multiple comparisons using the FDR procedure (Benjamini and Hochberg, 1995), accounting for our testing of five kinds of neuronal modulations (spatial target, place, memory, serial position, and heading). Specifically, for each of the five behavioral variables of interest, we applied an FDR-correction to the five p-values from the binomial tests. Next, for any behavioral variables where we observed a significant overall level of responsive cells, we then conducted a follow-up post-hoc test to identify whether the level of that cell was significant in a particular subregion. This follow-up test was a separate one-sided binomial test, which we also corrected using the FDR procedure across the three different subregions that we examined.

## Acknowledgements

We are grateful to the patients for participating in our study. We thank Sang Ah Lee and Shachar Maidenbaum for providing helpful feedback on the manuscript. This work was supported by NIH Grants R01-MH061975, R01-MH104606, S10-OD018211, T32-NS064928 and the National Science Foundation.

## Author Contributions

MT, JM, SEQ, and JJ designed the study. MT, SEQ, CSI, REG, JTW, EHS, SAS, CAS, MRS, AS,JMS, were involved in data collection. MT performed the data analyses. MT and JJ wrote the paper. All authors participated in editing and approved the final document.

## Declaration of Interests

The authors declare no competing interests.

## Supplemental Information

### Control for task version differences

We pooled the data from the two task versions because during the encoding portion of the tasks, which was the focus of this paper, subjects had similar behavioral and neural results. To confirm the neural result without pooling data, we checked that the proportions are similar when using only the object-cued version of the task. Excluding the data from the 3 subjects (4 sessions) who performed the location-cued version, there are 101 cells (instead of 131). The resulting proportions of cells modulated by each of the factors of interest from only the object-cued task version are described below, and are very similar to the results generated with data from both task versions.

19/101 of the cells were significantly modulated by the spatial target location, which is significantly above chance (*p* = 2.6 × 10^−6^; one-sided binomial test, FDR-corrected). The proportion of place-like cells was 7/101 (*p* = 0.24; one-sided binomial test, FDR-corrected). The proportion of heading-modulated cells was 11/101 (*p* = 0.02; one-sided binomial test, FDR-corrected), and there were 18/101 serial position cells (*p* = 6.5 × 10^−6^; one-sided binomial test, FDR-corrected). Finally, the number of memory-related cells actually was above chance at 10/101 (*p* = 0.038). Spatial-target, heading-modulated, and serial position cells all remain significant when calculated for this subset of the data, which supports our decision to pool the data from the two task versions. An interesting direction of future research would be to collect data from the same subjects performing both task versions and analyze to probe the degree to which task-related neurons may be modulated according to the different task demands.

### Interactions with neurons modulated by subsequent memory performance

To identify potential interactions between the spatial coding patterns we found, we tabulated the neurons that showed multiple behavioral modulations (Table S2). Of the 11 cells that showed significant memory-related firing-rate changes, six demonstrated memory-related increases while five showed decreases (see Figure S3c). Of the eleven memory-related cells, three were spatial-target cells, one was a heading-modulated cell, and none were place-like cells. Of the three cells that fulfilled the criteria for both memory and spatial-target cells, one showed a significant interaction between those two factors in a follow-up two-way ANOVA, which is notable but does not provide significant evidence for a link between these phenomena at the population level (*p* = 0.19, one-sided binomial test). Additionally, two serial position cells showed significant memory-related modulation, which was not a statistically significant overlap (p=0.89; *χ*^2^ test, df=1).

Notably, looking at firing rate by distance error for the object-cued task sessions gives very similar results. We find that 8 out of 101 cells exhibited a significant correlation between firing rate and distance error (*p* < 0.05). Six of those were also identified with our analysis of memory cells, which is a statistically significant correspondence (*p* = 2.4 × 10^−12^, *χ*^2^ (1) = 49.1). This similar result indicates that splitting the data into recalled and unrecalled captures a very similar pattern of results compared to measuring the raw error distance.

### Analysis of single cell counts with more stringent threshold

To confirm that our results are statistically robust on the single-cell level, we computed the number of significant cells for each parameter using a stricter p threshold of alpha = 0.01. Using this threshold and then a one-sided binomial tests with alpha = 0.01, we continued to find significant proportions of MTL cells responding to spatial target position (10*/*131, *p* = 2.4 × 10^−6^; one-sided binomial test, FDR-corrected), heading direction (5*/*131, *p* = 0.013; one-sided binomial test, FDR-corrected), and serial position (10*/*131, *p* = 2.4 × 10^−6^; one-sided binomial test, FDR-corrected). We also find a significant proportion of cells modulated by the subjects’ current position with this threshold (5*/*131, *p* = 0.013; one-sided binomial test, FDR-corrected). The proportion of memory-related cells remains below chance (1*/*131, *p* = 0.73; one-sided binomial test, FDR-corrected) By showing that our main results remain largely the same with a more stringent threshold, we demonstrate that individual cells show the effects we report strongly.

### Analysis for effects of epilepsy

To rule out the possibility that the effects we observed related to electrodes being located in abnormal brain tissue, we re-calculated our main results excluding units that were localized to what was clinically determined to be the seizure onset zone (SOZ). This included 15 units from three subjects (from Patients 2, 10, 12), leaving 116 units. Our main findings remained consistent after this exclusion. The proportion of spatial-target cells (23*/*116 = 20%), heading-modulated cells (14*/*116 = 12%), and serial position cells (17*/*116 = 15%) found in regions outside the SOZ all remained significantly greater than chance (*p* = 7 × 10^−8^, *p* = 0.0033, *p* = 1.7 × 10^−2^, respectively; one-sided binomial tests, FDR-corrected). The proportion of place-like cells, 6*/*116 = 5%, remained below chance (p = 0.53; one-sided binomial test, FDR-corrected), as did the proportion of memory-related cells (9*/*116 = 8%, *p* = 0.16; one-sided binomial test, FDR-corrected). The results we find are overall very similar to the original proportions. As such, we believe that the results are very unlikely to be due to pathological activity.

### Supplemental movie 1 caption

Example trial of the object-cued task, followed by an example trial of the location-cued task.

---

**Figure S1:**
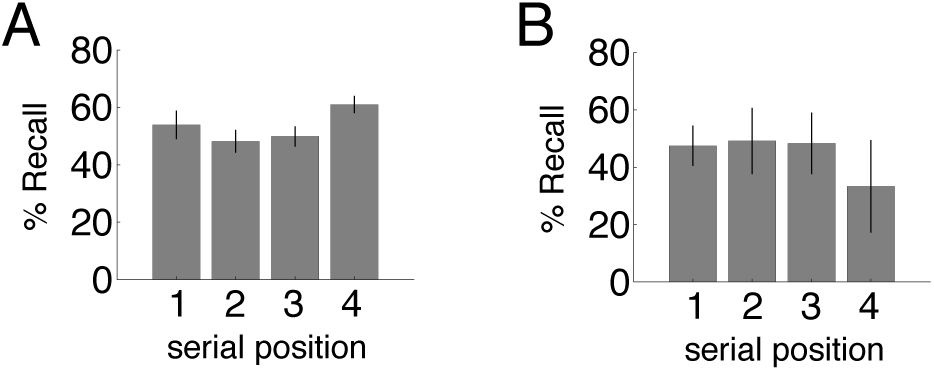
Recall by serial position. Mean and SEM of the percentage of object-location pairs successfully recalled, split by the serial position the object was encountered in, for the object-cued task sessions in panel A (p = 0.008; repeated measures ANOVA) and the location-cued task session in panel B (p = 0.1; repeated measures ANOVA).

**Figure S2:**
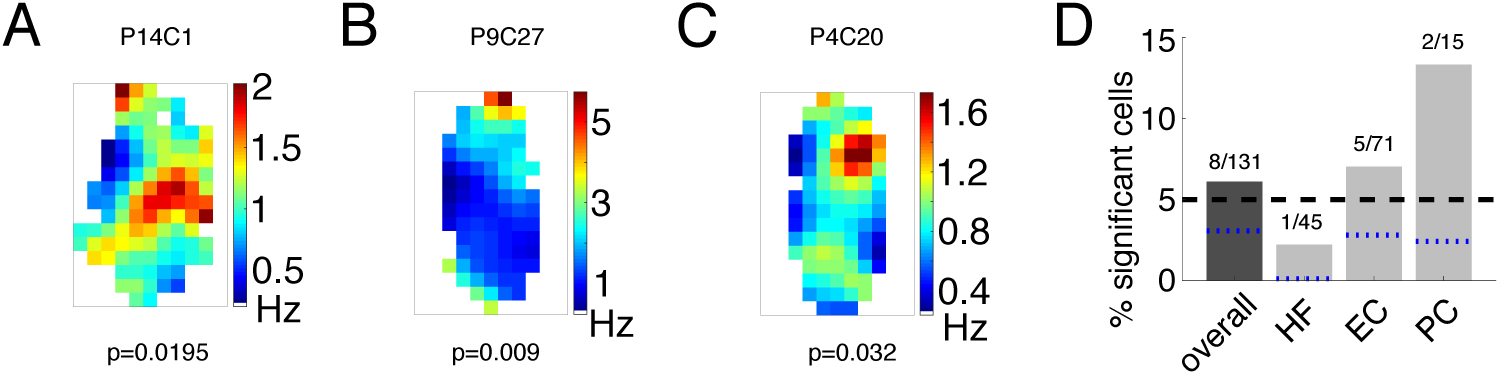
Neural activity related to the subject’s current location. A. Firing rate map of navigation activity binned by subject location for a right parahippocampal cortex neuron from Patient 14. This cell’s activity is significantly modulated by the subject’s virtual position (permutation-corrected ANOVA, p = 0.0195). B. Example neuron from left EC in Patient 9 significantly modulated by subject position (p = 0.009). C. Example neuron from Patient 4 in right EC significantly modulated by subject position. (p = 0.032). D) Percentage of significant spatially-tuned cells by region. The black dashed line represents the 5% false positive rate, and the blue dotted lines are the lower 95% confidence interval from the one-sided binomial test for each bar.

**Figure S3:**
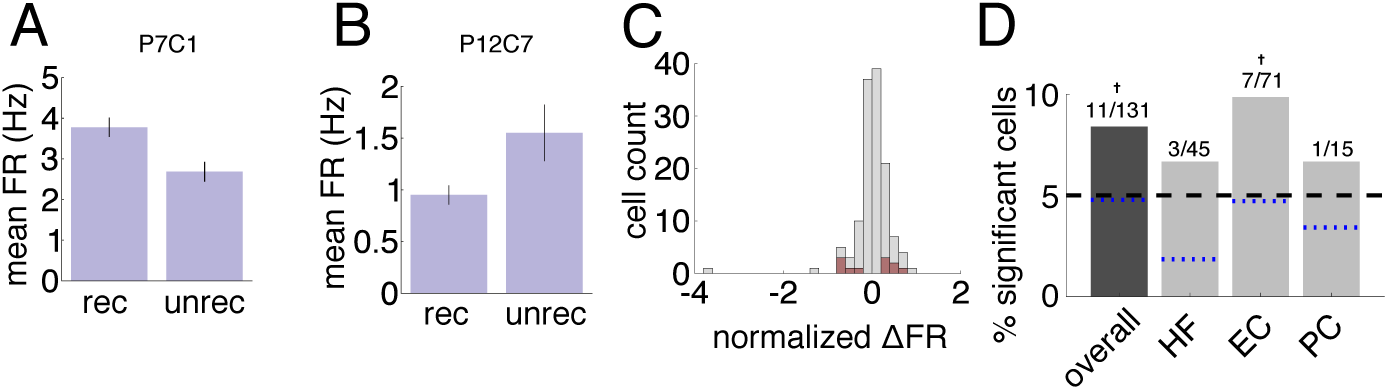
Neural activity related to subsequent memory. Firing rate by subsequent memory performance for two example cells significantly modulated by subsequent recall. Error bars are SEM of the trial means in each condition. A. Example neuron from Patient 7 in the left PRC, significantly modulated by subsequent memory (p = 0.017). B. Example neuron from Patient 12 in the right EC, p = 0.044. C. Histogram of cell count by normalized firing rate between conditions ((rec-unrec)/rec). Red indicates the significant cells. D. Percentage of significant memory-related cells by region. Symbols above the bars indicate p-values from a one-sided binomial test for each proportion (* * p < 0.01, *p < 0.05, +p < 0.1). The black dashed line represents the 5% false positive rate, and the blue dotted lines are the lower 95% confidence interval from the one-sided binomial test for each bar.

**Figure S4:**
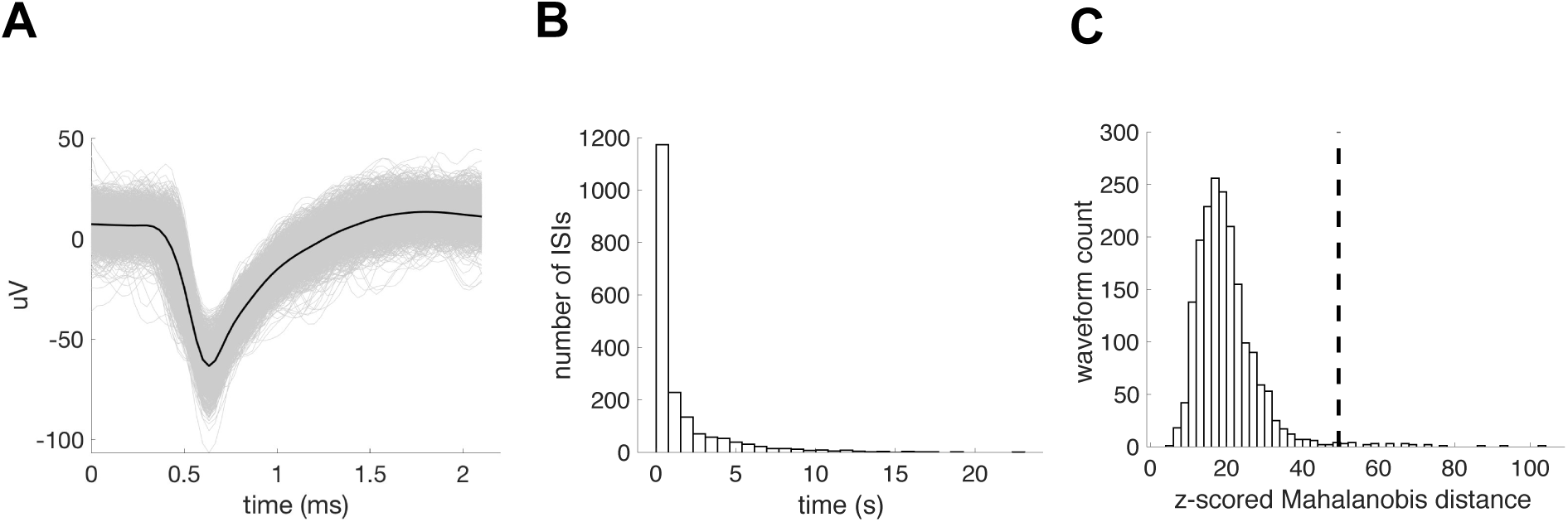
Single-unit classification example. Example unit from Patient 15. A. All waveforms across the task session in grey, and the average waveform in black. B. Histogram of interspike intervals (ISIs), showing less than 5% (0.3163%) contamination of the refractory period of 3ms. C. Histogram of normalized Mahalanobis distances from the mean of the cluster to each waveform. We compared that to a chi-squared distribution, and the dotted line represents the cutoff for the odds of there being any data this extreme less than 50%. Fewer than 5% of the waveforms (1.4%) were classified as outliers.

**Figure S5:**
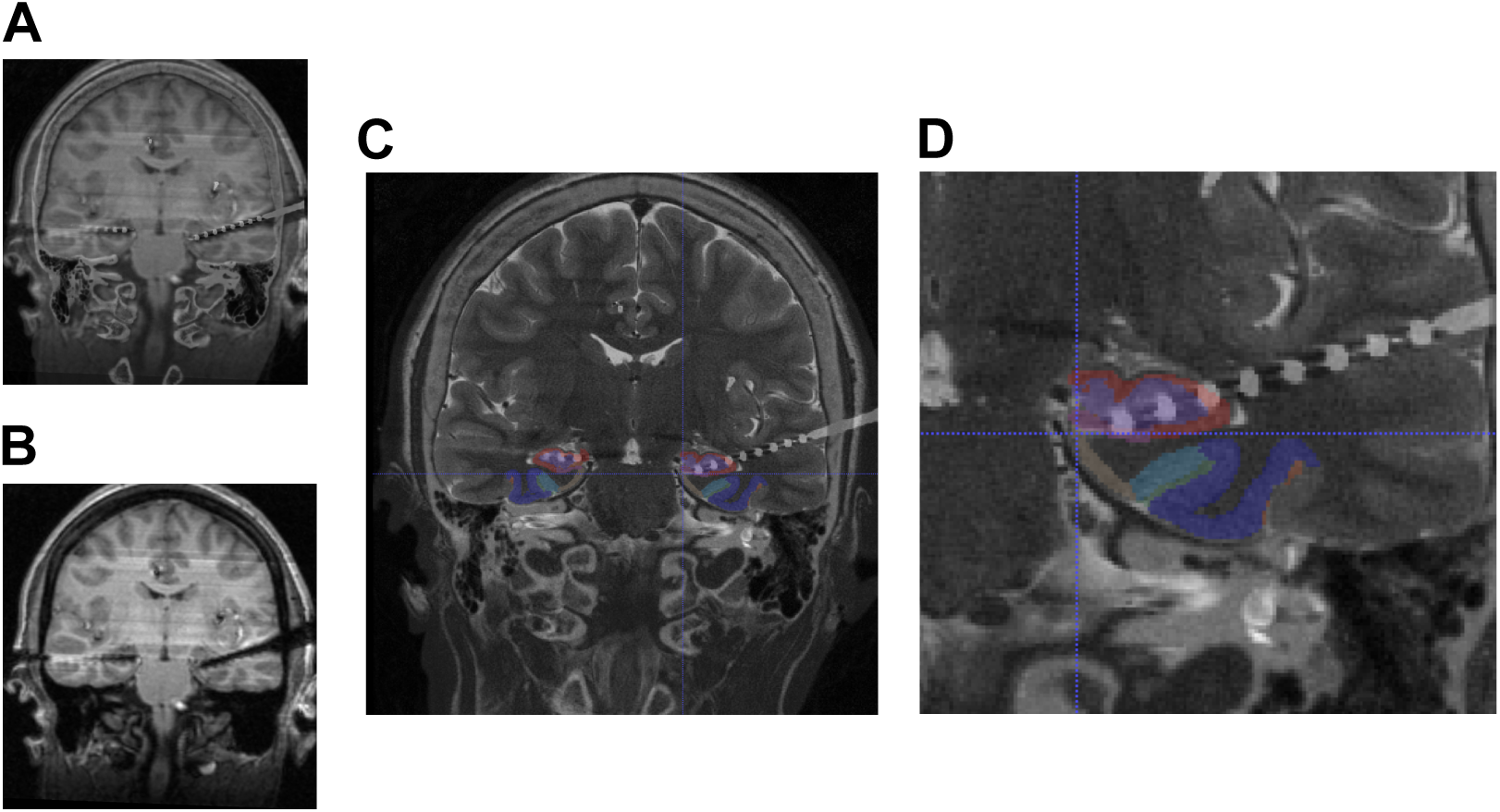
Example microwire localization images. Example localization of microwire bundle from Patient 11. A. Coregistered postoperative T1 MRI and CT scan. B. Postoperative T1 MRI scan without CT. C. Preoperative T2 structural MRI coregistered to postoperative CT. MTL subregion automatic segmentation is shown superimposed in color. ERC (tan), BA35 (light blue), BA36 (dark blue), subiculum (pink), dentate gyrus (purple), CA1 (red). D. Close-up of the same image as in C, with crosshair on microwire bundle, showing localization to ERC.

**Figure S6:**
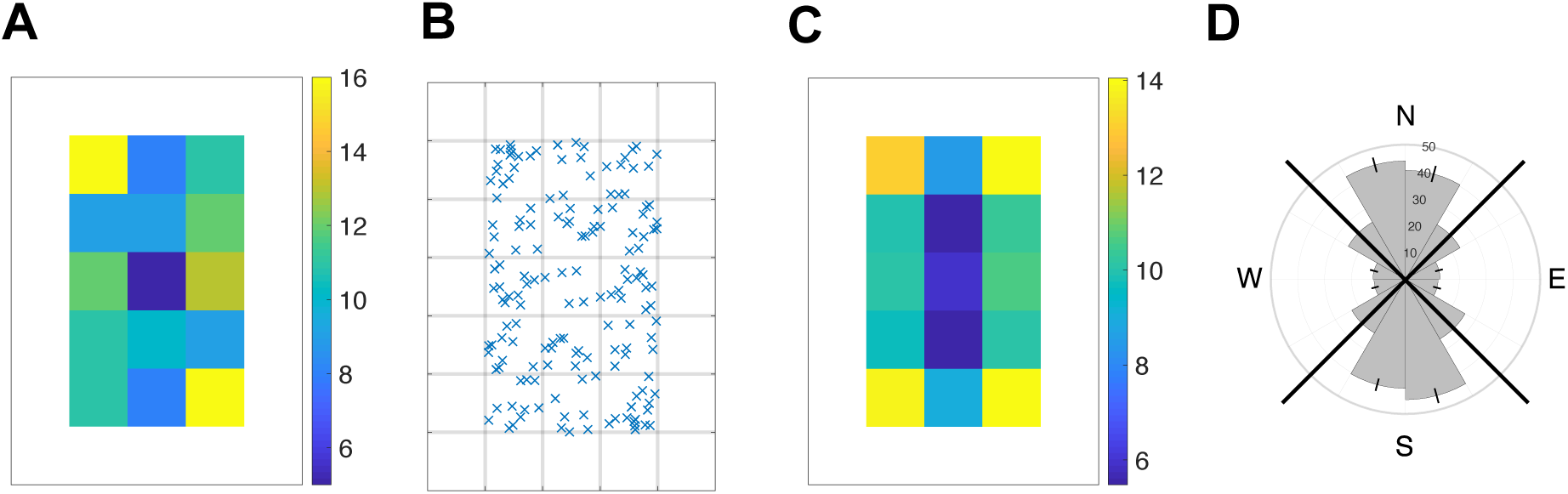
Distribution of chest position and heading direction across sessions. A. Number of chests in each location in a representative example session. Chests are never located in the outer border of the environment, shown in white. B. Same data with an ‘X’ indicating each chest location, with gray lines indicating the environmental binning. C. Number of chests in each grid location, averaged across all sessions. D. Circular histogram indicating the seconds (mean and SEM) spent in each heading direction during navigation, across all sessions. Black diagonal lines indicate NESW quadrants.

**Figure S7:**
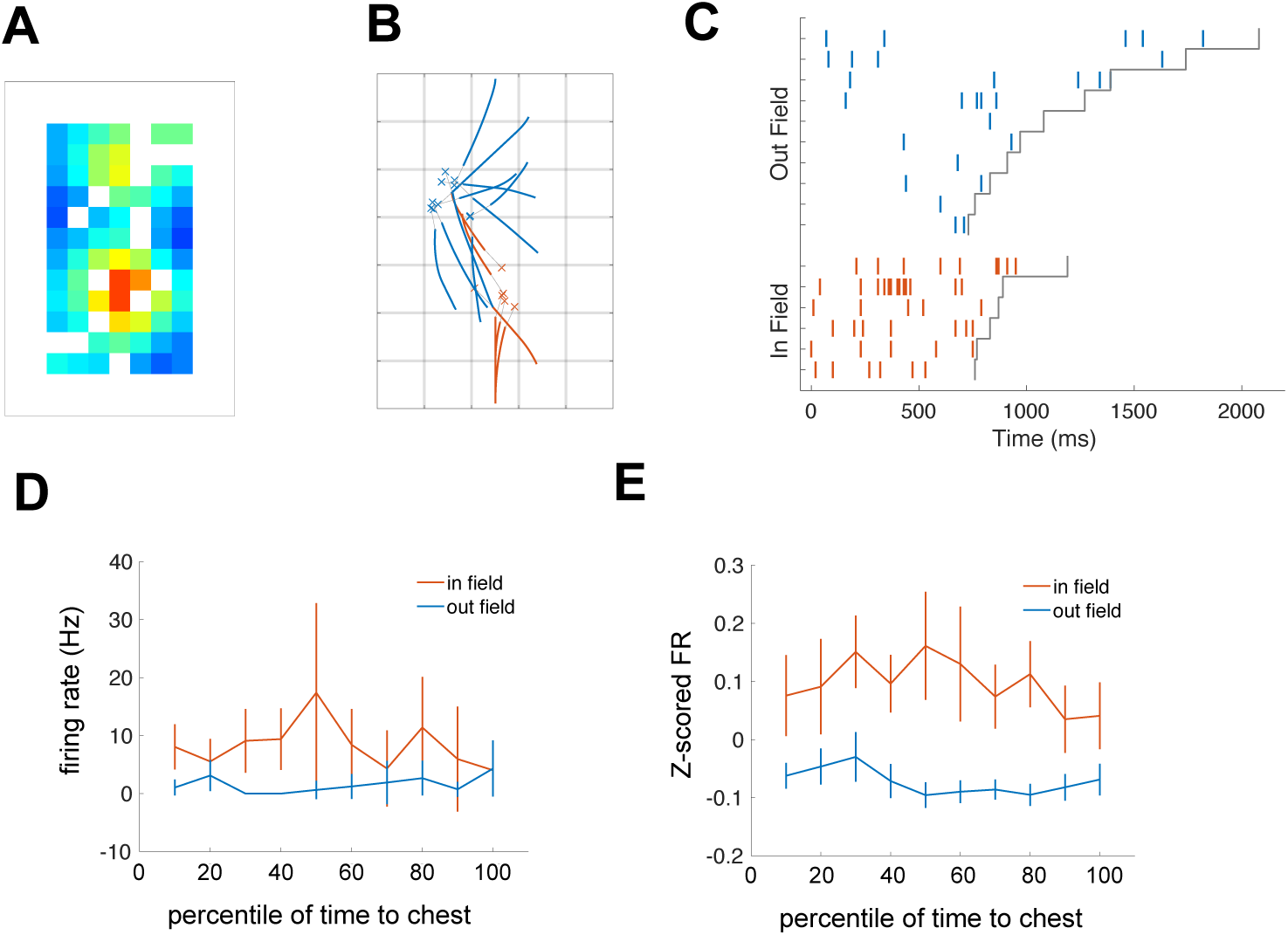
Spatial-target cell firing rates leading to chests in and out of the firing field. A. Firing rate map binned by chest location for example spatial target cell (Fig 2B in manuscript). B. Paths to chests in field and out of field for example cell. In field paths are in orange, out of field paths are in blue. C. Raster of spikes during the paths to in and out of the field for same example cell, again color coded by in and out of field, and sorted by navigation time for each of the fields. D. Mean and 95% confidence intervals of firing rates by normalized time to chest for example cell, split by in and out of field paths. E. Mean and 95% confidence intervals of the z-scored firing rates across all significant spatial-target cells.

**Figure S8:**
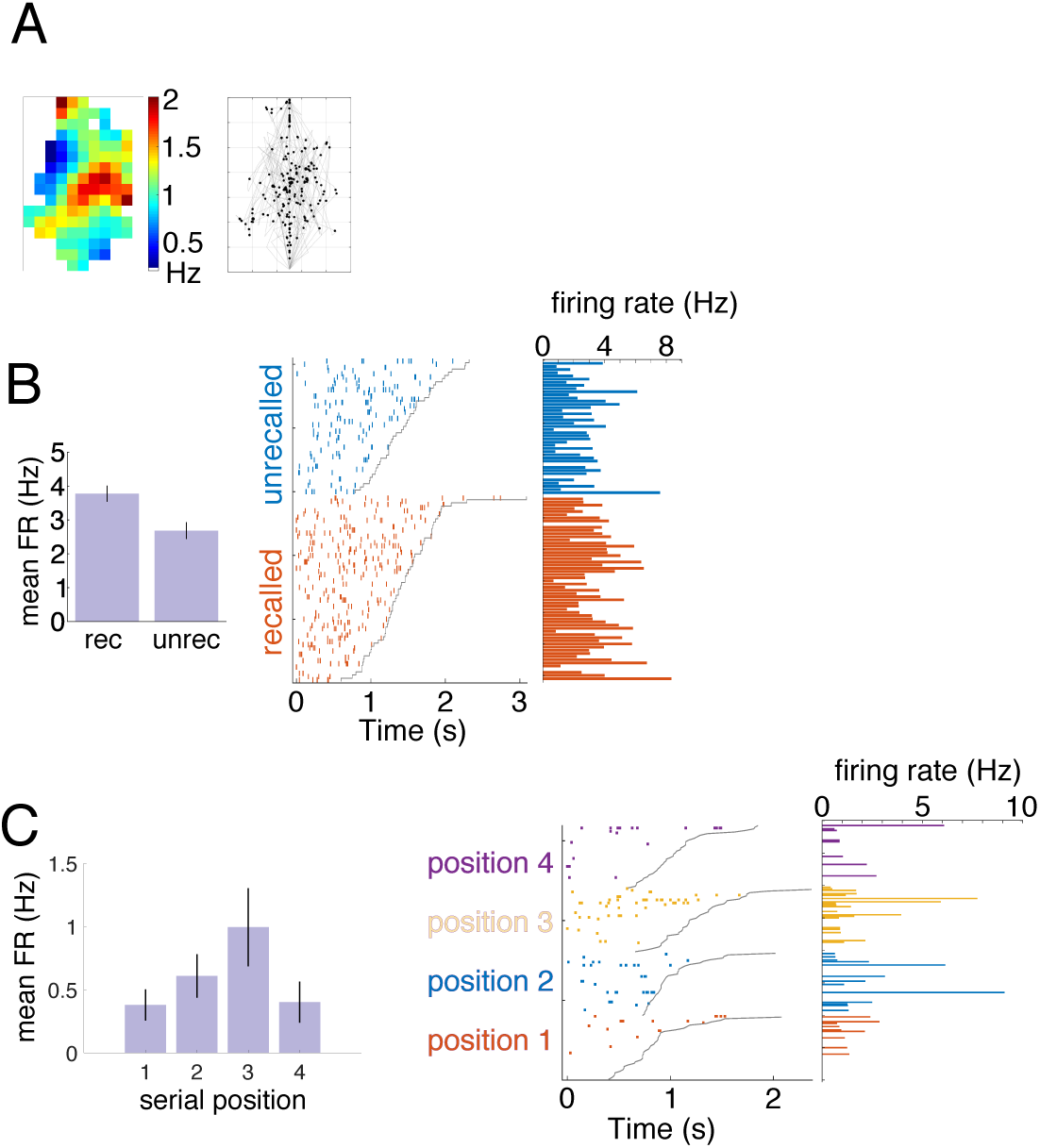
Example raster plots for place, memory, and serial position cells. A. Example place-like cell (p = 0.0195) in right PHC, from Patient 14 Cell 1 (same as in Figure S2a). Left plot shows firing rate map, and right panel shows the subject’s path in grey with a black dot indicating each spike. B. Example memory cell (p = 0.017) in left PRC, from Patient 7 Cell 1 (same as in Figure S3a). Left plot shows mean firing rate by recall. Right plot shows spike rasters and corresponding firing rates split by the two conditions, and ordered by path duration for each group. Grey line indicates when navigation ended. C. Example serial position cell (p = 0.025) in left EC, from Patient 9 Cell 17 (same as in Figure 4b. Left plot shows mean firing rate by each serial position. Right plot shows spike rasters and corresponding firing rates split by serial position, and ordered by path duration for each group. Grey line indicates when navigation ended.

**Figure S9:**
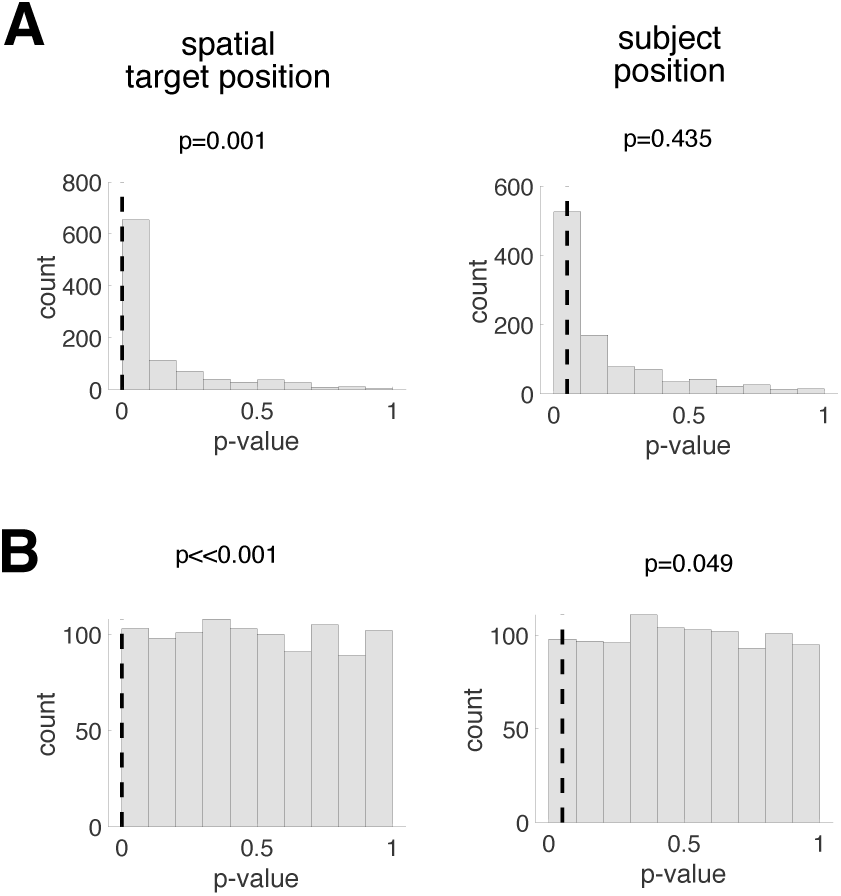
Comparison of p-value distributions generated from different shuffling procedures for example spatial-target cell. A. Histogram of p-values from ANOVA assessing a parameter’s modulation of firing rate for the observed data (dotted line) versus shuffled data (gray). Surrogate distribution is generated by circularly shifting spikes. Left plot is for spatial target position, and right plot is for subject position. This cell’s activity is significantly modulated by the spatial target position (p=0.001) but not subject position (p=0.435). B. Same plots as in A, but surrogate distributions are generated by randomly shuffling the spikes. Using this procedure both spatial target position and subject position are significant (p ≪ 0.001; p = 0.049, respectively).

**Figure S10:**
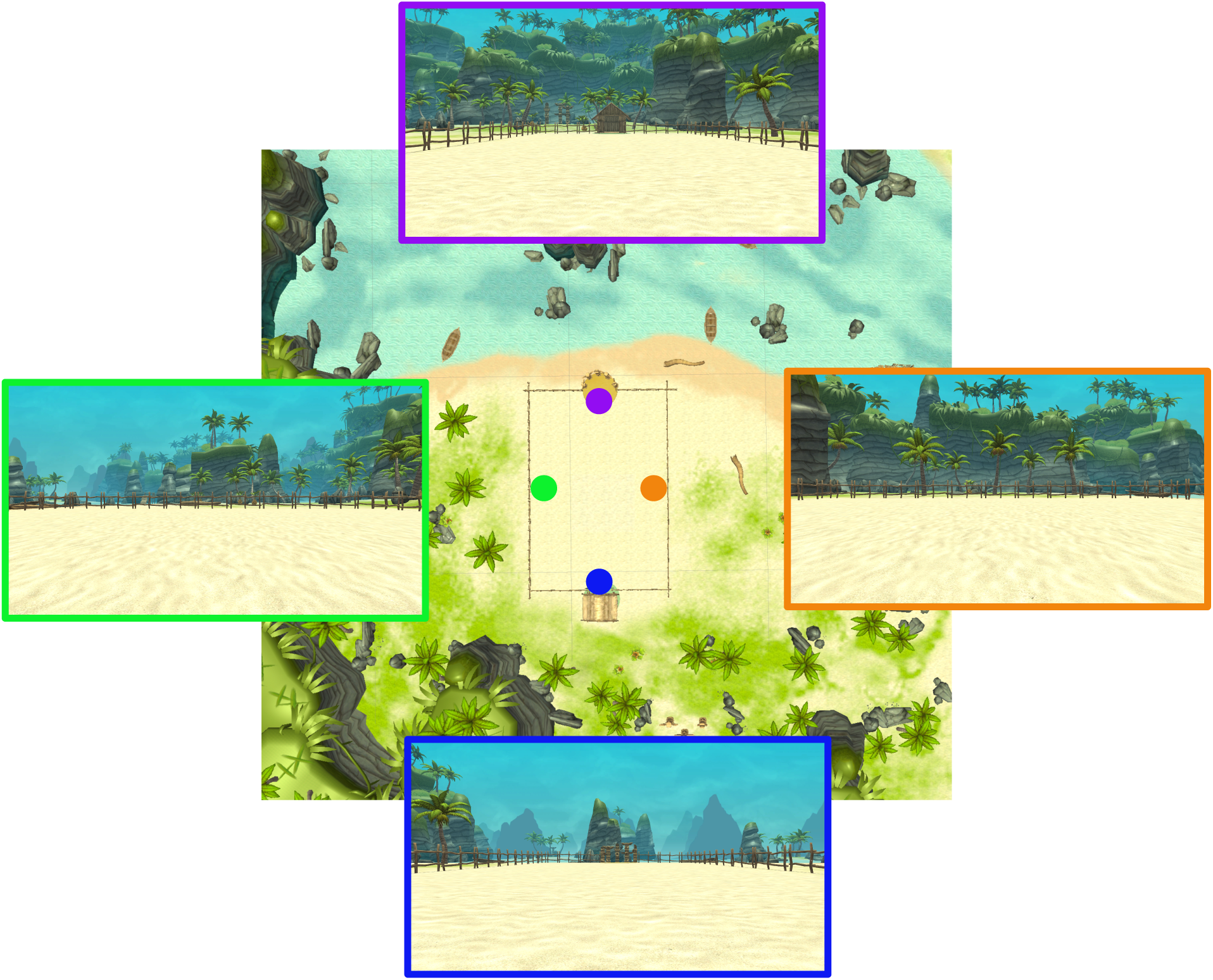
Treasure Hunt environment. Central image is a map (overhead view) of the Treasure Hunt environment. The rectangular arena that subjects navigate in is in the middle. At the top end of the arena there are totem poles in a semicircle, and water in the background behind them. At the bottom end of the arena there is a hut, and behind that there is a forest. The four overlayed panels show the subjects’ view from each of the locations indicated by the colored circles, with the border color matching the corresponding location.

**Table S1:**
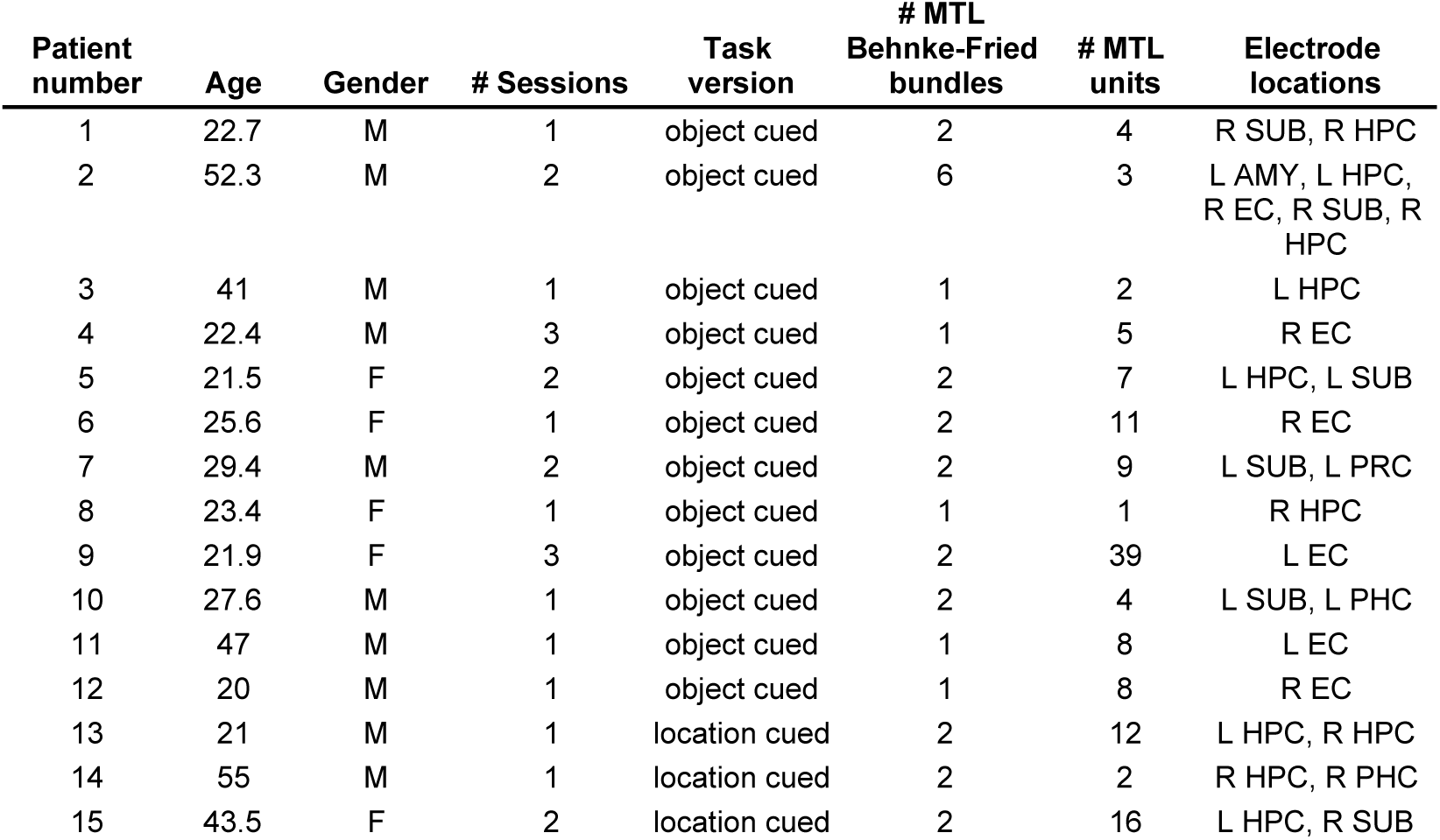
Patients and unit information. Table indicates each patient’s demographics and their MTL unit counts. R/L: right/left; HPC: hippocampus, SUB: subiculum, AMY: amygdala, EC: entorhinal cortex, PRC: perirhinal cortex, PHC: parahippocampal cortex.

**Table S2:** Significant cell counts. Number of cells with significant responses at different recording sites. The number of significant cells across MTL subregions are shown for each main effect and combination of main effects. Significance is determined using a shuffle-corrected ANOVA at alpha = 0.05. Counts of cells that were significant for place and another main effect, or or memory and another main effect, are not shown because place-like cells and memory cells were not found at significant proportions. Counts of cells that did not show any significant effects are also shown.

| Brain region | Total # units | Spatial target | Place | Heading | Memory | Serial position | Spatial target and place | Spatial target and heading | Spatial target and serial position | Heading and serial position | No effects |
| --- | --- | --- | --- | --- | --- | --- | --- | --- | --- | --- | --- |
| HF | 45 | 9 | 1 | 6 | 3 | 3 | 0 | 1 | 0 | 0 | 24 |
| EC | 71 | 12 | 5 | 7 | 7 | 16 | 0 | 1 | 3 | 4 | 37 |
| PC | 15 | 5 | 2 | 3 | 1 | 1 | 0 | 1 | 0 | 1 | 5 |
| **total** | **131** | **26** | **8** | **16** | **11** | **20** | **0** | **3** | **3** | **5** | **66** |

